# Closing the Loop on Phage-bacteria Coevolution

**DOI:** 10.64898/2025.12.15.694372

**Authors:** Joshua Pearson, Kirill Sechkar, Harrison Steel

## Abstract

Bacteria and their viruses, bacteriophages (phages), continually coevolve in nature. In contrast, laboratory-based coevolution experiments usually last less than a month before becoming dormant or extinct as one species is outcompeted by the other. Consequently, there is a poor understanding of phage-bacteria coevolution and hence the long-term efficacy of bacteriophage therapies (an approach to tackling antimicrobial resistance). We propose a novel approach to coevolution experiments that would address this challenge: instead of open-loop resource-constrained cultures, we develop a closed-loop control approach to stabilise the typically unstable or oscillatory phage-bacteria population dynamics. Achieving this requires the control system to compensate for delays in phage incubation and respond to an evolving system, while only measuring bacterial density. To this end, we develop a model of phage-bacteria dynamics, prototype delay-compensating predictive control strategies, and demonstrate a measurement-aware state observer. Overall, this approach shows the ability to stabilise coevolution, avoiding the common outcomes of unstable dynamics or winner-takes-all competition.

## 1. INTRODUCTION

Control theory applied to Engineering Biology has predominantly focussed on ‘rational’ design of gene circuits, yet it equally promises to advance evolutionary approaches, such as Adaptive Laboratory Evolution. In ALE, the problem shifts from directly designing a high-performance strain to engineering the selective pressures that drive its emergence. Control techniques have already been applied to evolution to understand drug-resistance in cancers (Fischer et al. (2015)) and the emergence of antimicrobial-resistant bacteria (Toprak et al. (2013)). In this paper, we consider a related challenge: controlling the coevolution of bacteria and their viruses, bacteriophages (phages), in a bioreactor.

Bacteriophages only infect specific bacteria, leaving human cells unaffected, and hence are attractive agents for treating bacterial infections. As a therapy, phages are more selective than antibiotics, allowing treatments to be tailored to target individual bacterial species. Unfortunately, as with antibiotics, bacteria can evolve resistance to phages, for example by modifying or even removing the surface receptors used by phages for binding (Hasan and Ahn (2022)). However, a key advantage of phages (unlike antibiotics) is that they can evolve to overcome resistance, often by changing their spike proteins (see Figure 1). Bacteria and bacteriophages have sufficiently short lifecycles, and sufficiently high selection pressure, that the two can co-evolve on the timescales of laboratory experiments. For the last 50 years, phage-bacteria coevolution experiments have used an open-loop approach (Chao et al. (1977)). However, these experiments suffer from large ecological oscillations and usually end after a month, with either phage or bacteria out-competing the other and driving it to extinction or dormancy (Dennehy (2012)).

**Fig. 1.**
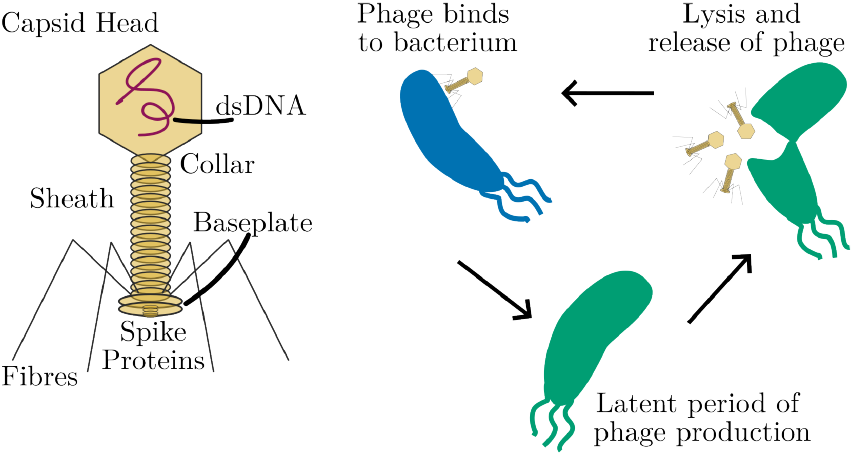
Left: diagram of Bacteriophage T4, showing capsid head containing double-stranded DNA and spikes that bind to *E. coli* receptors. Right: the life cycle of a lytic phage, requiring a host bacterium for replication.

To address this challenge, we propose a novel control-enabled approach for coevolution experiments. Our approach involves growing bacteria at regulated low densities, setting up a density-dependent competition where an equilibrium in ecological dynamics exists. To replicate, phages must collide with and bind to bacteria, making the rate of replication proportional to the product of bacterial concentration and phage concentration; in contrast, for a fixed nutrient availability, bacteria reproduce at a fixed growth rate per cell, irrespective of concentration. This difference in density-dependence of growth presents a promising avenue for controlling phage and bacterial populations. We hypothesise that these dynamics can be stabilised by controlling media turnover in a bioreactor (removal of media and both species, and replacement with fresh media) to regulate ecological dynamics around an equilibrium point. This paper develops this idea, identifies conditions required for this fixed point to exist, and shows in simulation that it can be stabilised.

Technologies to measure changing phage concentrations in real-time and *in situ* do not exist, and hence we need to use the observed bacterial dynamics to infer phage concentrations. A further complication is that, because the system is evolving, the parameters of the ecological model will be changing over time. However, we address this by using a multi-rate observer for parameter-state estimation.

The purpose of these simulations is to prototype and de-risk experiments; a key question is determining design and implementation requirements for a bioreactor platform. For example, by determining appropriate phage and bacterial density ranges, we can outline sensor requirements (e.g. sensitivity and noise performance).

More widely, the monitoring and control of bioreactors are established applications of control theory. Our specific application has similarities with this broad family of control problems in bioprocesses: the system has uncertain, time-varying and non-linear dynamics, and key parameters cannot be directly measured, requiring inference. However, this bioreactor problem differs from most since it is not constrained by process scale-up; laboratory evolution only ever needs to be at millilitre scale.

To our knowledge, this is the first study to consider closed-loop control for phage coevolution. As this novel experimental method is developed in practice, it will generate highly valuable insights into the long-term behaviours of phages, with applications such as combating antimicrobial resistance. In this and other evolutionary design procedures, control engineering accelerates experiments, and biotechnological progress more broadly.

## 2. PROBLEM DEFINITION

In the proposed experiment, phages and bacteria are co-cultured in a bioreactor, and actuation is provided by varying the rate of replacement of cell media. The population dynamics model is given in Equations (1)–(6), with terms defined below. Values for the model lytic system of *E. coli* K-12 and Escherichia bacteriophage T4 were used in simulations, and are given in Table 1. Below, we discuss the modelling assumptions giving rise to these equations, and how they differ from the canonical model (Levin et al. (1977)). All simulations are available at https://github.com/j-c-pearson/closed-loop-coevolution.

**Table 1.** Parameters used in simulations.

| Phage parameters |  |  |
| --- | --- | --- |
| $B$ | 98 | Levin et al. (1977) |
| $\delta$ | $3.12 \times 10^{-8} [\text{mL hr}^{-1}]$ | Levin et al. (1977) |
| $\tau$ | 0.5 [hr] | Rabinovitch et al. (1999) |
| *E. coli* parameters |  |  |
| $\mu$ | 1.8 [$\text{hr}^{-1}$] | Corrao et al. (2023) |

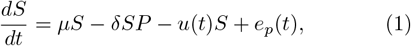

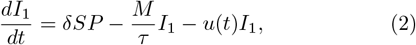

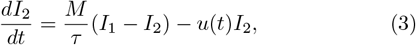

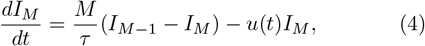

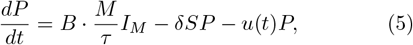

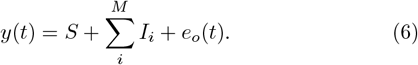

Our system variables are:

- *S* is the concentration of susceptible bacteria [mL^−1^],
- *I*_*i*_ is the concentration of infected bacteria, in pseudostate *i* [mL^−1^],
- *P* is the concentration of phages [mL^−1^],
- *u*(*t*) is the rate of media turnover [hr^−1^],
- *y*(*t*) is the observation of optical density [mL^−1^],
- *e*_*p*_(*t*) *~ N* (0, *σ*_*p*_) is the process noise,
- *e*_*o*_(*t*) *~ N* (0, *σ*_*o*_) is the observation noise.

For a susceptible cell to become infected, a phage must collide with it and bind successfully, which can be modelled using mass-action-like dynamics (Levin et al. (1977)), such that the rate is proportional to the concentration of both species and a binding constant *δ*. The production of phages then occurs within the cell for a latent period *τ* before lysis (destruction of the cell membrane) takes place. We have modelled this delay using the ‘linear chain trick’ (Smith (2011)), where the delay is broken into a series of *M* pseudostates, *I*_1_, *I*_2_, …, *I*_*M*_. We used *M* = 5, which has been found to be appropriate for similar models (Geng et al. (2024)). When lysis occurs, the bacterium dies and the burst number (*B*) of phages is released. These phages then go on to bind to other bacteria, continuing the replication cycle (Figure 1).

The susceptible (uninfected) bacterial cells are modelled as growing in exponential phase with growth rate *µ*. Previous phage coevolution studies have used open-loop chemostats (Levin et al. (1977)) or batch cultures (Geng et al. (2024)), where resources are expected to limit populations. However, we propose to operate at densities orders of magnitude lower to control ecological dynamics (see below), where bacterial populations will not be additionally resource-constrained, and hence resource limitation is not considered in our model.

The emergence of new phage and bacterial strains through mutation (and hence coevolution) can profoundly impact control system performance. For instance, if the emergence of a partially-resistant bacterial strain is not detected, the controller may remove media faster than the phages can replicate, driving the phage populations extinct. To investigate our state-estimation algorithm’s performance in such a circumstance, we model mutation by introducing new subpopulations, which have the same dynamics but potentially different parameters, partway through the simulation. Upon detection of a new strain, the system can be rescued by changing the controller’s parameters or mixing in fresh, non-resistant bacteria, although this remedial action is not simulated in this work.

## 3. CONTROL APPROACH

We hypothesise that a key reason for *evolutionary* out-competition in laboratory experiments is the oscillatory *ecological* dynamics, which provide very strong selection. We propose that, by stabilising the ecological dynamics, we can *unnaturally* decrease selection and prevent out-competition.

### 3.1 Turnover rate actuates ecological dynamics

Controlling bacterial concentration, and hence the encounter rate of phages and bacteria, can stabilise the ecological dynamics at a fixed point. Qualitatively, the growth of bacteria is proportional to the number of bacteria, whereas the proliferation of phages is proportional to the product of the number of bacteria and the number of phages. As such, the dilution of the bioreactor with fresh media, which acts to proportionately diminish both organisms’ counts, is an asymmetric actuator; at low bacterial densities, phages are washed out (i.e. without first finding bacterial targets), while at high bacterial densities, the phage concentrations spike. Thus, around an equilibrium point, increasing dilution rate *u*(*t*) would compensate for too high phage concentrations, while decreasing *u*(*t*) would do the opposite.

To determine an operating point, we use the delay differential equations (not the ODE approximation given above):

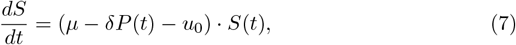

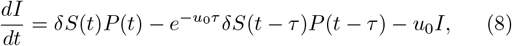

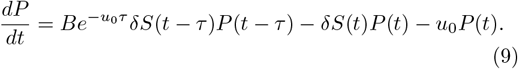

Since, at a fixed point, *S*^*∗*^ = *S*(*t −τ*) = *S*(*t*) etc., Equations (7), (8), (9) can be solved, giving closed form solutions in terms of parameters *B, δ, τ, µ, u*_0_:

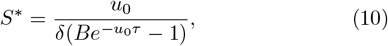

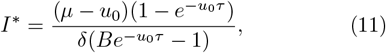

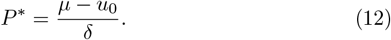

The open-loop (error-free) rate of media turnover, *u*_0_, is a free parameter that needs to be selected. For it to be a non-trivial equilibrium, *u*_0_ = *µ −δP*\*, and so there is an upper limit of *u*_0_ *< µ* provided by the condition *P*\* *>* 0. The equilibrium concentration of bacteria increases with *u*_0_, while the concentration of phages decreases. Higher bacterial concentrations should improve the signal-to-noise ratio of the sensors. However, *u*_0_ should be sufficiently far from *µ* (since *u*_0_ = *µ* gives *P*\* = 0) that phage concentrations are robust to process noise. To balance these considerations, we picked *u*_0_ = 1.25, which corresponds to a fixed point of *S*^*∗*^ = 8.1 *×* 10^5^, *I*^*∗*^ = 1.6 *×* 10^5^, *P*\* = 1.9 *×* 10^7^ cells ml^−1^ for the default parameters of the system, and can be re-estimated when new parameters are detected. Previous coevolution experiments used bacterial densities of *~* 10^7^*−* 10^8^ cells ml^−1^ (OD_600_ *~* 0.01 *−*0.1) (Levin et al. (1977)). At these densities, no fixed point exists (which we prove in Appendix A). We note that operating at our proposed densities may have been infeasible previously, but bioreactors introduced in Corrao et al. (2023) include high-resolution OD sensing, making this practical; see the noise discussion below.

Using Lyapunov’s indirect method, we can assess the stability of our operating point. The linearised system has two unstable eigenvectors, but these correspond to perturbations to bacterial concentration, which can be controlled; the phage concentration (which is inferred but never explicitly controlled) is locally stable. Therefore, closed-loop control is required to stabilise the ecology around this equilibrium.

### 3.2 Control system design

A key characteristic of our design problem is the presence of delays in phage incubation. To compensate for these, we use a Smith Predictor architecture for our controller, which compares model predictions, both before and after the effect of delay, to the observed signals, as shown in Figure This error signal is then fed into a low-level controller (*C* in Figure 2); a PI controller performed well here. Our approach deviated slightly from the architecture described in Normey-Rico and Camacho (2007) since, instead of a delay between input and state, the delay exists in the autonomous dynamics. Therefore, the inputs affect the state throughout the prediction horizon and need to be specified; using the target (error-free) input of *u*_0_ = 1.25 proved stable and effective.

**Fig. 2.**
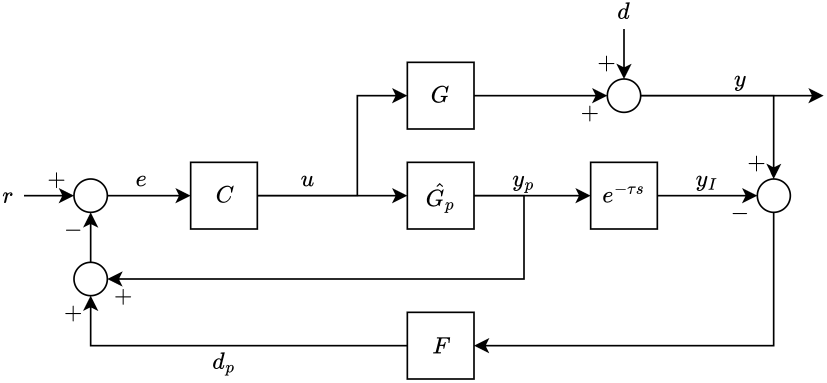
The Smith Predictor uses a model (*Ĝ*_*p*_) to make a prediction (*y*_*p*_) for the future output. This is added to the filtered difference (*d*_*p*_) between the previous prediction for that time (*y*_*I*_) and the measurement (*y*), providing feedback. The error signal (*e*) is then passed to a low-level controller (*C*) to create the actuation signal (*u*).

The observer needs to track both species’ concentrations and system parameters, while only observing the bacterial concentration. A measurement of the bioreactor’s optical density (proportional to *y* in Equation (6)) measures the concentration of all (susceptible and infected) bacteria, but cannot detect phage, since they are below the diffraction limit of the wavelength used. Instead, our approach infers phage concentrations from observed bacterial dynamics (phage predation acts as a time-varying deficit in growth rate). Tracking system parameters is important to prevent the system from destabilising, since bacterial resistance (lowering *δ*) can lead to phages being washed out.

Parameter-state estimation can be achieved by both joint (augmented state) and dual (bootstrapping) approaches — see Wan and van der Merwe (2000) for an overview. Both were trialled, and a dual approach showed superior performance. In addition to a lower computational cost, dual parameter-state estimation allows timescales to be separated. By tracking parameters less frequently, more informative samples are used, improving numerical properties, while still re-estimating at a faster frequency than the system changes. In our implementation, the state estimate is updated every step, using the latest parameter estimates, whereas the parameters are updated every *n*_*Dual*_ = 10 steps, using the new state estimates.

For the underlying filter, an Unscented Kalman Filter (UKF) is used for both parameters and states. In simulation, the UKF had comparably accurate estimates to an Extended Kalman Filter (EKF) but succeeded in propagating uncertainty through the nonlinearity. In contrast, the EKF’s estimates became overconfident with time, which is unacceptable for an (ideally) indefinite experiment. As discussed in Wan and van der Merwe (2004), the effect of sigma-point sampling parameters *α, β, κ* on the UKF is only partially understood and can significantly affect performance. In general, *α* controls the spread of sigma points, *κ* provides secondary control scaling, often used to account for dimensionality, while *β* can encode prior understanding of the distribution, with *β* = 2 optimal for a Gaussian distribution. Using parameters recommended in Wan and van der Merwe (2000) (*α* = 10^*−*3^, *β* = 2, *κ* = 0), the state covariance matrix **P**_*xx*_ became non-positive definite, a problem noted in Julier and Uhlmann (1997). Wan and van der Merwe (2004) note that parameter estimates are much more sensitive to *α, β, κ* than state estimates. By instead using parameters implicitly used in Julier and Uhlmann (1997) (*α* = 1, *β* = 0, *κ* = *−*1), a stable filter was achieved, although the covariance estimate will only be accurate to second order.

As a strain with higher fitness is fixed in the population, the ‘bulk’ parameters are expected to interpolate between those of the old strain and the new strain. This can be modelled naturally using correlated/filtered noise, achieved by augmenting parameter estimates with a parameter momentum estimate, as discussed in Simon (2006). (We note that, in the general case, it is *not* true that one-strain approximations of parameters are concentration-weighted averages. However, if strains emerge and are fixed sequentially, we expect to observe approximately sigmoidal transitions between old and new parameters.) The momentum approach showed significant improvement over the most common approach, a persistence mechanism, where parameters drift under white noise (Simon (2006)). Persistence mechanisms functioned, but struggled to distinguish changes in species concentration from changes in parameters. Therefore, incorporating prior understanding of how parameters will evolve allows us to create a better filter.

## 4. SIMULATION RESULTS

We first simulated our control system’s efficacy in situations where no new strains were emerging. The left side of Figure 3 shows both the ‘true’ values in the simulation (solid lines) and the observer’s estimate (dashed lines), showing close agreement. For comparison, a simulation of open-loop behaviour is shown on the right side of Figure 3, showing the system entering unstable oscillations. In this and all other simulations, we introduce sensor noise *σ*_*o*_ and process noise *σ*_*p*_ (set to 3 *×* 10^3^ and 6 *×* 10^3^ [ml^−1^ hr^−1^], respectively), based on estimates from our lab’s bioreactors. That the system is stabilised in the presence of noise provides initial proof of concept that the phage-bacteria ecology can be controlled.

**Fig. 3.**
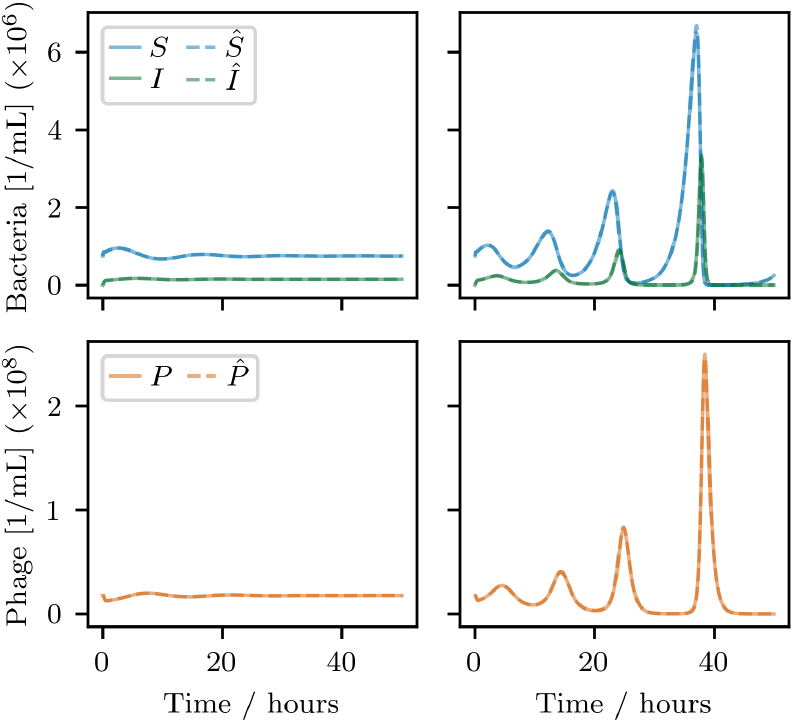
Left: Ecological dynamics are stabilised by our control system, in the case of fixed parameters. Right: open-loop behaviour (*u*_0_ = 1.25) for comparison. Here, and in the graphs below, hatted variables are the observer estimates, and are plotted with dashed lines.

When experimentally realised, in our biological system the strongest selection is expected to be on the phage binding parameter, *δ*, since this determines whether the phage replicates (and hence whether a bacterium is lysed or not). Therefore, we simulated new bacterial and phage strains emerging as subpopulations with different *δ* parameters (an increase corresponding to more infective phages, and a decrease to more resistant bacteria). To allow the strains to fully emerge, these simulations were run for 250 hours, instead of the 50 hours used in other simulations. Figure 4 shows the system successfully controlling the ecological dynamics as a fitter phage strain emerges. We had similarly good performance in simulations of a more resistant bacterial strain emerging (see Figure 5 in Appendix C).

**Fig. 4.**
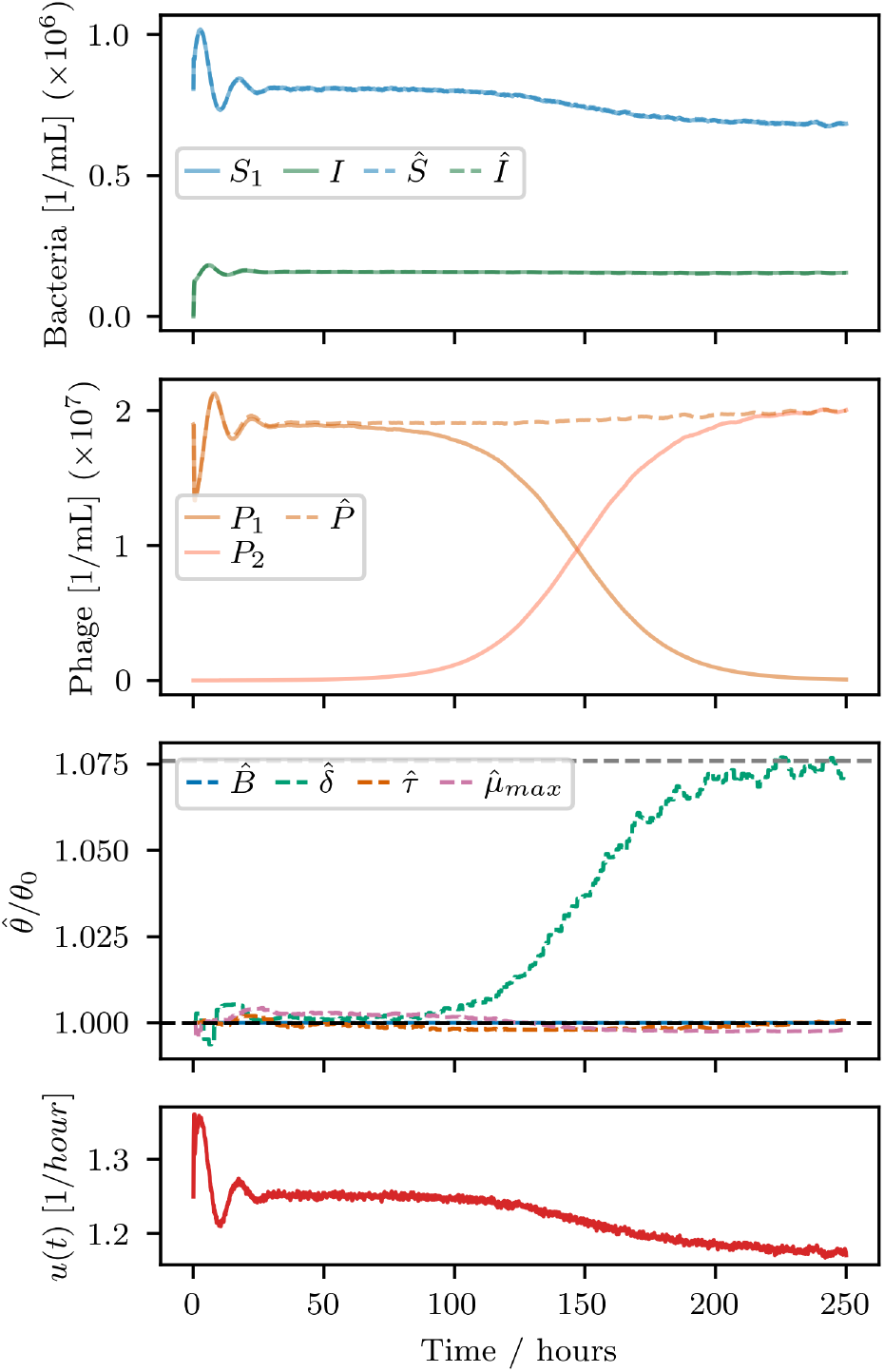
Simulation of second phage strain *P*_2_ with higher binding constant *δ* emerging and fixing. *δ*_1_ = 2.9 *×* 10^*−*8^ [mL hr^−1^], *δ*_2_ = 3.12 *×*10^*−*8^ [mL hr^−1^], giving a 7.6% increase (marked by the grey dashed line). The observer closely tracks the change in *δ*, estimating a 7.3% increase by the end of the simulation.

**Fig. 5.**
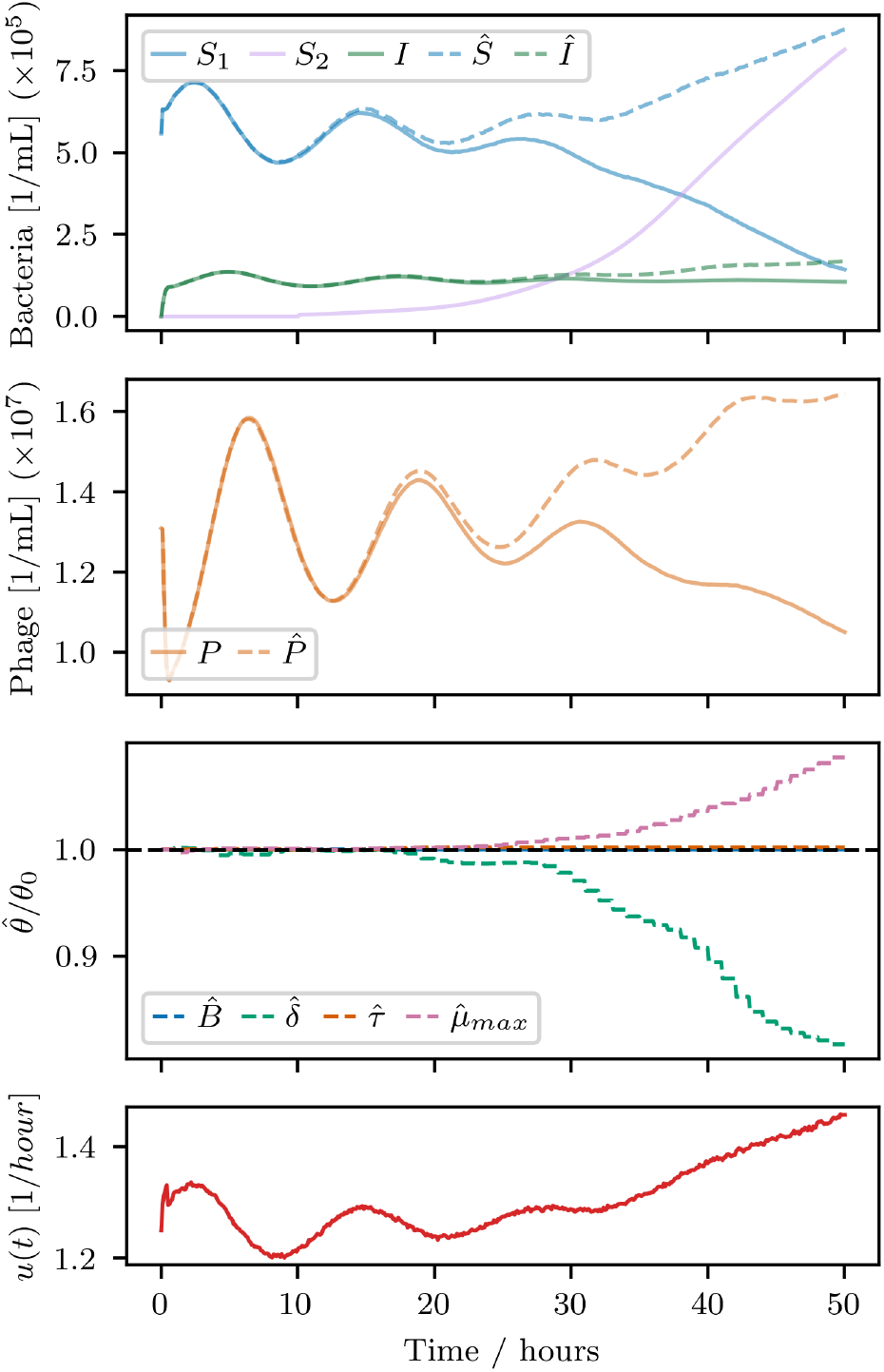
Simulation of a semi-resistant bacterial strain *S*_2_ emerging and fixing. The bacterial strain has such a competitive advantage that it rapidly fixes and the observer destabilises, with 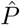 diverging from *P*. *δ*_1_ = 4.2 *×* 10^*−*8^ [mL hr^−1^], *δ*_2_ = 2.9 *×* 10^*−*8^ [mL hr^−1^], giving a 31% decrease in *δ* for *S*_2_.

Since Bayesian filters have been used, specifying the prior understanding of parameter evolution (via the process noise covariance matrix *Q*_*param*_) is crucial. If *Q*_*BB*_ and *Q*_*ττ*_ (the variances of *B* and *τ*, encoded as diagonal elements in *Q*_*param*_) are too large (if the normalised standard deviation is more than *≈* 1%), then the observer cannot track the parameter changes, for example incorrectly attributing changes in *δ* to a change in *τ*. If *B* and *τ* are found to be evolving, this will pose a tough parameter-state estimation challenge, since there will exist a set of nearly indistinguishable solutions (with changes in both parameters and phage concentrations). In this situation, a particle filtering approach may be appropriate to model the multimodal distributions with cross-state dependence. In these simulations, some changes in growth rate were allowed, since mutations for resistance often carry a growthrate burden (Brockhurst and Koskella (2013)); when the low *δ* bacterial strain rapidly emerges in Figure 5, this is incorrectly attributed to a change in growth rate as well as a decrease in *δ*.

Although the model used by the observer is locally structurally identifiable, verified using STRIKE-GOLDD software (Villaverde et al. (2016)), this is insufficient for identification in practice (Wieland et al. (2021)). Structural identifiability analysis considers both noise-free and infinite amounts of (persistently exciting) data. However, in this system, parameters can evolve and so are not fixed over long time periods. Therefore, sufficient information must be acquired faster than the parameters change. As such, the observer fails if a new strain with vastly greater fitness rapidly emerges, as demonstrated in Figure 5.

While our system performed well under our estimates of noise levels, we wanted to understand whether the feasibility of the experiment was sensitive to these estimates. We therefore ran further simulations to understand the impact of actuation and sensor noise magnitude, shown in Figures C.2 and C.3 (available in Appendix sec:figures). While our proposed control system was fairly robust to sensor noise, our state estimates diverged under the highest level of process noise tested, highlighting the need for careful tuning, and potentially development, of the dilution action when implementing these experiments.

## 5. CONCLUSION

We have proposed a closed-loop approach to bioreactor coevolution experiments, which could prevent evolutionary ‘dead-ends’ and enable long-term study of the coevolution of bacteria and phages. We have shown that there exists an unstable fixed point in the modelled system, which can be stabilised by control of the dilution rate. We then presented proof of concept for a control strategy to run these experiments, and simulations to explore their sensitivity, providing cautious optimism that our approach is feasible.

Our analyses have considered the paradigm of selective sweeps, where a strain with a beneficial mutation fixes in the population. Selective sweeps enable rapid adaptation, and are a common modelling choice for phage-bacteria coevolution (Brockhurst et al. (2014)). Developing simulations under the alternative paradigm of clonal interference (in which beneficial mutations compete with each other) would be an important step to further inform and de-risk laboratory implementations of this approach.

Identification remains challenging, particularly when multiple parameters change simultaneously. If both *δ* and *µ* vary significantly, then the observer may not be able to distinguish between a change in growth rate and simultaneous changes in phage concentration and *δ*. Further, if bulk values of *B* and *τ* are found to vary significantly over the experiment, then any point-estimate approach is unlikely to track the system; joint particle filtering approaches are an attractive solution in this scenario.

We hope this study will be of interest to those modelling and designing state estimators for bioreactors. More broadly, we have aimed to show how the intersection of control theory and biological modelling can inspire new experimental approaches. Such experiments have the potential to advance evolutionary approaches in Engineering Biology and to address fundamental scientific challenges, including the fight against antimicrobial resistance.

## ACKNOWLEDGEMENTS

The authors thank Antonis Papachristodoulou and Ting An Lee for insightful discussions throughout the work.

## DECLARATION OF GENERATIVE AI AND AI-ASSISTED TECHNOLOGIES IN THE WRITING PROCESS

During the editing of this work, the authors used Claude to improve clarity. After using this tool, the authors reviewed and edited the content, and take full responsibility for the content of the publication.

## Appendix A. PROOF OF NON-EXISTENCE OF FIXED POINTS AT PREVIOUSLY ATTEMPTED EXPERIMENTAL DENSITIES

Previous experiments have operated at *S ≈* 1 *×* 10^7^. We can prove that the uncontrolled dynamics (of an *E. Coli* and T4 system) do not have any fixed points at concentrations this high (without introducing resource-limiting conditions, as done in Chao et al. (1977)).

**Proof**. No fixed point point exists for *S >* 1.5 *×* 10^6^ cells mL^−1^, for any *u*_0_ *∈* R_*≥*0_

From the requirement that *P*\* *>* 0, Equation (12) introduces the upper bound *u*_0_ *< µ* for a fixed point. We can then maximise *S*^*∗*^ for 0 *< u*_0_ *< µ*.

Firstly, the derivative of *S*^*∗*^ with respect to *u*_0_:

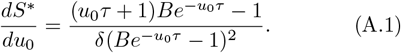

We can define the domain where 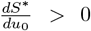 as *u*_0_ *∈* (*<u>u</u>, ū*), where *<u>u</u>, ū ≈ −* 2.0, 13.2 are the solutions to (*uτ* + 1)*Be*^*−uτ*^ = 1. Since ū *> µ* and *<u>u</u> <* 0, *S*^*∗*^ is monotonic as a function of *u*_0_ over the domain of valid *u*_0_ *∈* (0, *µ*).

Therefore, the largest susceptible bacterial density, *S*^*∗*^ occurs in the limit *u*_0_*→ µ*, giving 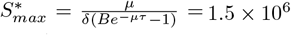 cells mL^−1^ (for *E. Coli* and T4). □

We note that introducing resource-limited conditions nullifies this result (Chao et al. (1977)), but we believe introduces a strong and unwanted evolutionary pressure for glucose-utilisation. Finally, we note that this is not a bifurcation (mathematically) but the movement of the fixed point to (biologically meaningless) negative values.

## Appendix B NOTES ON MODEL USED

In addition to binding to susceptible cells (*δSP* in Equation (1)), phages can bind to infected cells, potentially altering the dynamics (Geng et al. (2024)). This is included in some models as the Multiplicity of Infection (MOI, the number of phages binding to a bacterium), and is important when a large proportion of bacteria are infected. However, the proposed control strategy should prevent the infected cells from reaching a high enough density for MOI’s effects to be significant, and so it is not modelled in this work (we verified this in additional simulations, the code for which is available at https://github.com/j-c-pearson/closed-loop-coevolution).

## Appendix C ADDITIONAL FIGURES

**Fig. C.1.**
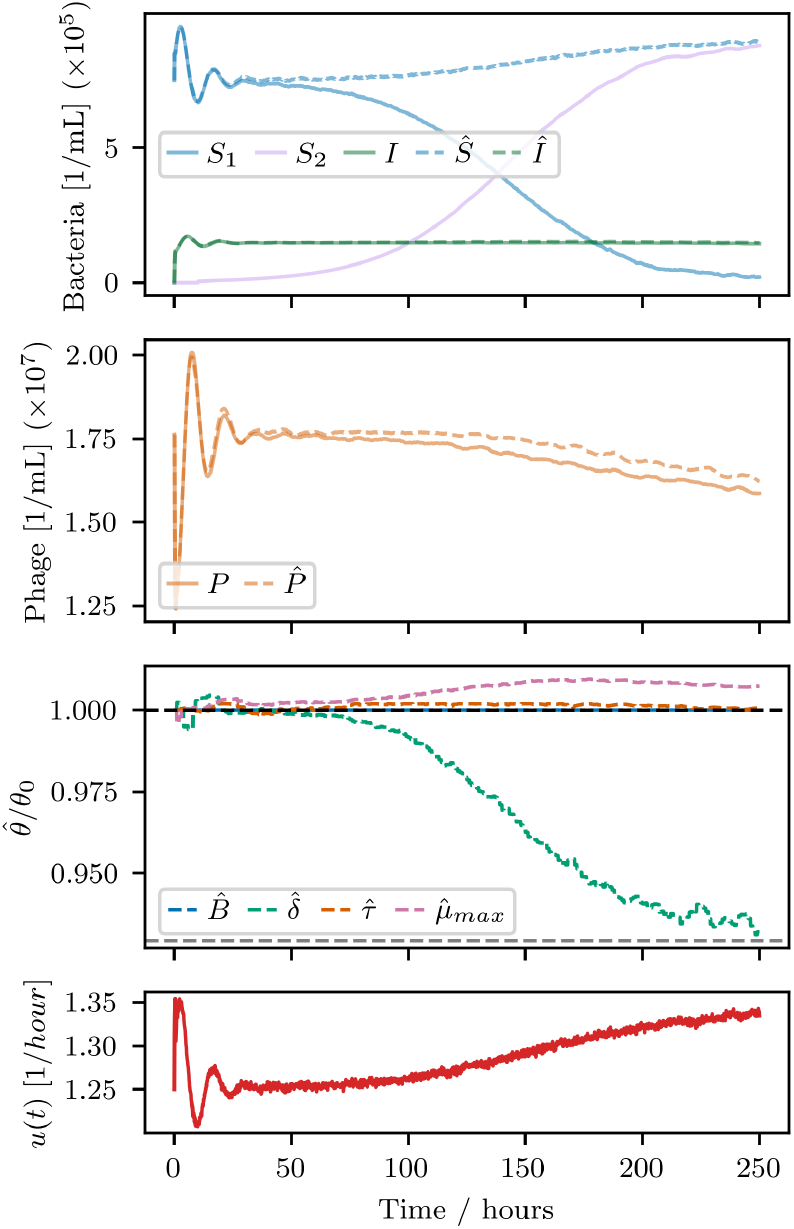
Simulation of second bacterial strain *S*_2_ with lower binding constant *δ* emerging and fixing. *δ*_1_ = 3.12 *×* 10^*−*8^ [mL hr^−1^], *δ*_2_ = 2.9 *×*10^*−*8^ [mL hr^−1^], giving a 7.1% decrease (marked by the grey dashed line). The observer tracks the change in *δ*, estimating a 6.8% decrease by the end of the simulation. Although the phage concentration estimate diverges slightly at *~* 100 hours, it then re-converges.

**Fig. C.2.**
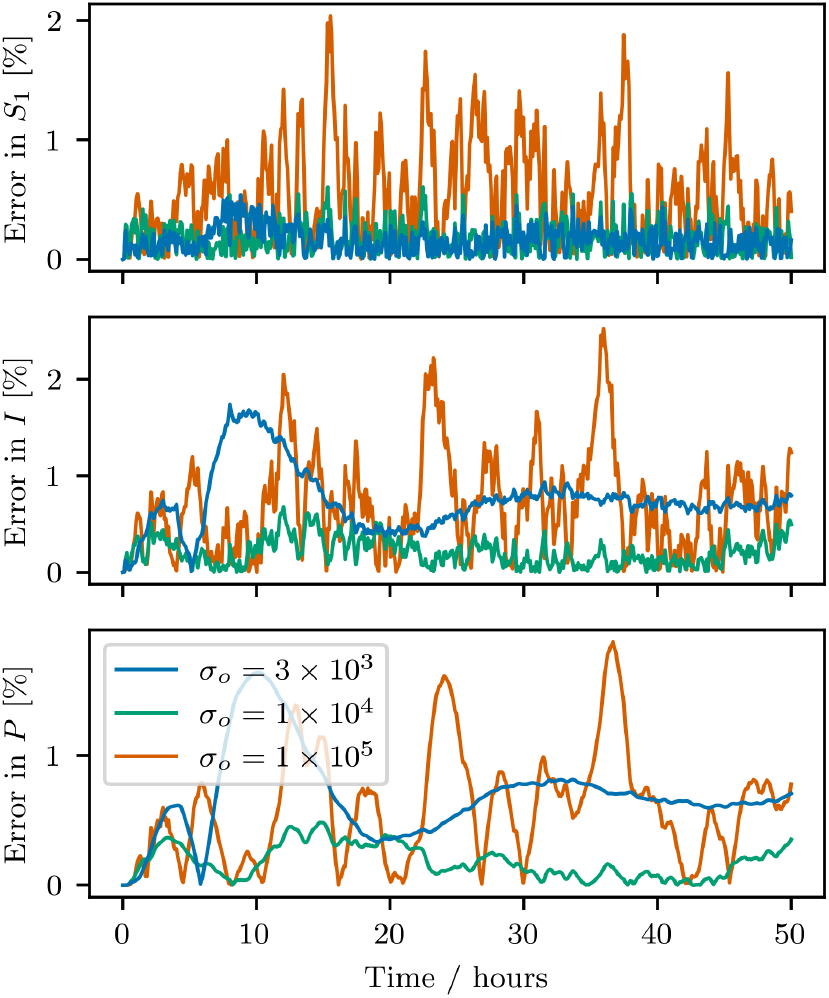
Effect of sensor noise on percentage error in species estimates. The observed state (*S*) unsurprisingly has lower error for a given noise level. Error calculated as 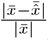, where 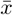 is the average true value over 50 runs, and 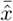 is the average estimate of the 50 runs. Units of noise are cells / (mL hour)

**Fig. C.3.**
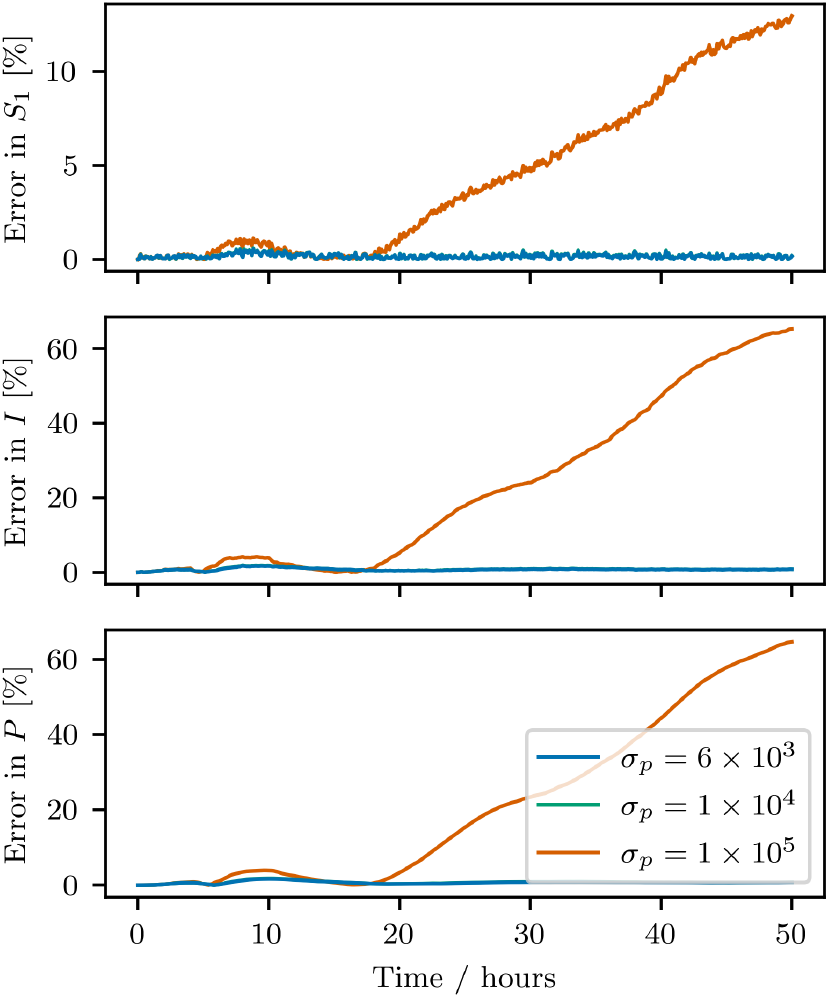
Effect of process noise on percentage error in species estimates. Units of noise are cells / (mL hour).

## REFERENCES

Brockhurst, M.A., Chapman, T., King, K.C., Mank, J.E., Paterson, S., and Hurst, G.D. (2014). Running with the red queen: the role of biotic conflicts in evolution. Proceedings of the Royal Society B: Biological Sciences, 281(1797).

Brockhurst, M.A. and Koskella, B. (2013). Experimental coevolution of species interactions. Trends in ecology & evolution, 28(6), 367–375.

Chao, L., Levin, B.R., and Stewart, F.M. (1977). A com-plex community in a simple habitat: an experimental study with bacteria and phage. Ecology, 58(2), 369–378.

Corrao, M., Vigouroux, A., and Steel, H. (2023). An automated platform for accelerating adaptive laboratory evolution. In Proceedings of the 2023 International Workshop on Bio-Design Automation.

Dennehy, J.J. (2012). What can phages tell us about host-pathogen coevolution? International journal of evolutionary biology, 2012(1).

Fischer, A., Vázquez-García, I., and Mustonen, V. (2015). The value of monitoring to control evolving populations. Proceedings of the National Academy of Sciences, 112(4), 1007–1012.

Geng, Y., Nguyen, T.V.P., Homaee, E., and Golding, I. (2024). Using bacterial population dynamics to count phages and their lysogens. Nature Communications, 15(1), 7814.

Hasan, M. and Ahn, J. (2022). Evolutionary dynamics between phages and bacteria as a possible approach for designing effective phage therapies against antibiotic-resistant bacteria. Antibiotics, 11(7), 915.

Julier, S.J. and Uhlmann, J.K. (1997). New extension of the Kalman filter to nonlinear systems. In Signal processing, sensor fusion, and target recognition VI, volume 3068, 182–193. Spie.

Levin, B.R., Stewart, F.M., and Chao, L. (1977). Resource-limited growth, competition, and predation: a model and experimental studies with bacteria and bacteriophage. The American Naturalist, 111(977), 3–24.

Normey-Rico, J. and Camacho, E. (2007). Control of deadtime processes. Springer.

Rabinovitch, A., Hadas, H., Einav, M., Melamed, Z., and Zaritsky, A. (1999). Model for bacteriophage T4 development in Escherichia coli. Journal of Bacteriology, 181(5), 1677–1683.

Simon, D. (2006). Optimal state estimation: Kalman, H infinity, and nonlinear approaches. John Wiley & Sons.

Smith, H. (2011). An introduction to delay differential equations with applications to the life sciences. Springer.

Toprak, E., Veres, A., Yildiz, S., Pedraza, J.M., Chait, R., Paulsson, J., and Kishony, R. (2013). Building a morbidostat: an automated continuous-culture device for studying bacterial drug resistance under dynamically sustained drug inhibition. Nature protocols, 8(3), 555– 567.

Villaverde, A.F., Barreiro, A., and Papachristodoulou, A. (2016). Structural identifiability of dynamic systems biology models. PLoS computational biology, 12(10).

Wan, E.A. and van der Merwe, R. (2000). The unscented Kalman filter for nonlinear estimation. In Proceedings of the IEEE 2000 adaptive systems for signal processing, communications, and control symposium.

Wan, E.A. and van der Merwe, R. (2004). The unscented Kalman filter. In S. Haykin (ed.), Kalman Filtering and Neural Networks. John Wiley & Sons.

Wieland, F.G., Hauber, A.L., Rosenblatt, M., Tönsing, C., and Timmer, J. (2021). On structural and practical identifiability. Current Opinion in Systems Biology, 25, 60–69.

